# Local field potentials and single unit dynamics in motor cortex of unconstrained macaques during different behavioral states

**DOI:** 10.1101/2022.04.28.489967

**Authors:** Richy Yun, Irene Rembado, Steve I. Perlmutter, Rajesh P. N. Rao, Eberhard E. Fetz

**Affiliations:** Department of Bioengineering, Science and Engineering, University of Washington; Department of Physiology and Biophysics, Science and Engineering, University of Washington; Department of Allen School for Computer, Science and Engineering, University of Washington; Center for Neurotechnology, University of Washington; Washington National Primate Research Center, University of Washington; Allen Institute for Brain Science

## Abstract

Different sleep states have been shown to be vital for a variety of brain function, including learning, memory, and skill consolidation. However, our understanding of neural dynamics during sleep and the role of prominent LFP frequency bands remain incomplete. To elucidate such dynamics and changes between different behavioral states we collected multichannel LFP and spike data in primary motor cortex of unconstrained macaques for up to 24 hours using the Neurochip3. Each 8 second bin of time was classified into awake and moving (Move), awake and at rest (Rest), REM sleep (REM), or non-REM sleep (NREM) by using dimensionality reduction and clustering on the average spectral density and the acceleration of the head. LFP power showed high delta during NREM, high theta during REM, and high beta when the animal was awake. Cross-frequency phase-amplitude coupling typically showed higher coupling for deeper sleep between all pairs of frequency bands. Two notable exceptions were high delta-high gamma and theta-high gamma coupling during Move, and high theta-beta coupling during REM. Sorted single units showed decreased firing rate with deeper sleep, though with higher “bursty” patterns during NREM compared to other states. Spike-LFP synchrony showed high delta synchrony during Move, and higher coupling with all other frequency bands with deeper sleep.

These results altogether are consistent with previous findings showing reactivation of cortical circuitry during sleep, which may be moderated by delta band LFP.

## Introduction

The discovery of rapid-eye movement (REM) sleep by Aserinski and Kleitman showed the presence of active brain processes during sleep, which catalyzed studies on different sleep stages (Aserinsky & Kleitman, 1953). Findings have shown REM sleep to play a role in regulating the autonomic nervous system activity and to be linked with spatial and emotional memory consolidation (Boyce et al., 2016; Rechtschaffen, 1998; Siegel, 2005). Similarly, non-REM (NREM) sleep has been shown to be tied to skill consolidation and memory formation (Ramanathan et al., 2015; Siclari & Tononi, 2017).

Studies have furthered our understanding of different sleep stages by exploring the dynamics of local field potential (LFP) frequency band power and single-unit spiking characteristics in both the cortex and various deep brain structures during sleep. Slow waves and delta frequency band are typically present across the brain during NREM sleep, and high theta power in the hippocampus is present during REM sleep (Rechtschaffen & Kales, 1968; Silber et al., 2007). Single units show changes in firing rate as well as firing patterns during sleep, due to changes in connectivity and excitability, potentially reflecting reactivations of relevant cortical circuit patterns to solidify learning and memory (Arbune et al., 2020; Ramanathan et al., 2015; Tononi & Cirelli, 2014; Xu et al., 2019). Several other features of cortical activity, such as k-complexes and sleep spindles during NREM sleep, have also been identified to be associated with different stages of sleep (Fernandez & Lüthi, 2020; Huber et al., 2004; Ulrich, 2016).

Additional findings have shown that certain plasticity mechanisms, such as long-term potentiation/depression and synaptic tagging and capture, occur exclusively during sleep (Kim et al., 2005; Vecsey et al., 2018). Subsequent research has applied this knowledge by enhancing slow waves prevalent during NREM sleep (Bellesi et al., 2014; Marshall et al., 2006) or delivering stimulation locked to delta to augment learning and memory (Rembado et al., 2017). However, despite our expanding tools and techniques to take advantage of sleep mechanisms, the neural dynamics underlying sleep remain unclear.

Various measures of local field potential (LFP) coupling and spike-LFP synchrony have been commonly used to elucidate mechanisms of neural dynamics. Cross-frequency phase-amplitude coupling of different LFP frequency bands, thought to reflect communication between brain circuitry, has been shown to be modulated by task performance, cognitive engagement, and memory formation in various brain regions (Canolty & Knight, 2010; Jensen & Colgin, 2007). Single units are strongly synchronized to specific phases of LFP frequency bands, notably to beta and gamma cycles in the motor cortex, suggesting beta to be a resting rhythm during movement execution and gamma to reflect local population activity (Buzsáki et al., 2012; Buzsáki & Wang, 2012; Engel & Fries, 2010; Murthy & Fetz, 1996a, 1996b). LFP and unit analyses have given us significant insight into neural signaling and the role of both spikes and LFPs in brain function, but there has yet to be a comprehensive analysis of cross-frequency coupling and spike-LFP synchrony for specific cortical sites during sleep and wake.

This study analyzes LFP and single unit dynamics in the macaque motor cortex during different behavioral states to better understand the mechanisms operating during different sleep stages as well as the roles of various LFP frequency bands. We first used the power spectral density of LFPs to distinguish between four major behavioral states shown to be relevant to plasticity and learning in the motor cortex: 1. awake and moving (Move), 2. awake and at rest (Rest), 3. rapid-eye movement (REM) sleep, and 4. non-REM (NREM) sleep. We tracked single-unit activity concurrently with LFP recordings and focused on 6 different frequency bands commonly observed in the cortex – delta, theta, alpha, beta, low gamma, and high gamma. Finally, we assessed state-dependent changes in cross-frequency coupling between pairs of the frequency bands as well as spike-LFP synchrony in each frequency band.

## Materials and Methods

### Experimental design

#### Implants and surgical procedures

The experiments were conducted on two male *Macaca nemestrina* monkeys (Monkeys J and K). Surgeries were performed under isoflurane anesthesia and aseptic conditions to implant multi-electrode Utah arrays. All procedures conformed to the National Institutes of Health *Guide for the Care and Use of Laboratory Animals* and were approved by the University of Washington Institutional Animal Care and Use Committee.

Implantation of the Utah array was guided by stereotaxic coordinates. A 1.5 cm wide square craniotomy was performed over the hand region of the primary motor cortex to expose the dura. Three sides of the exposed dura were cut to expose the cortex; a Utah array was implanted and the dura was sutured over the array. Two reference wires were inserted below the dura and two were inserted between the dura and the skull. The bone flap from the craniotomy was replaced and held in place by a titanium strap screwed onto the skull with 2.5 mm x 6 mm titanium skull screws. A second smaller titanium strap was fastened to the skull to secure the wire bundle. The connector pedestal for the array was attached to the skull with eight titanium skull screws and the incision closed around the pedestal base. Additional skull screws were placed around the base of the connector pedestal and a thin coat of dental acrylic (methyl methacrylate) was applied to the skull between the screws and the connector base for additional stability.

The animals also received a “halo” implant made with 3/8” aluminum forming an egg-shaped oval 17 cm long and 15.3 cm wide. Four titanium straps were affixed to the skull by titanium bone screws. Two were implanted bilaterally over the occipital ridge, and two were placed temporally bilaterally. After the plates integrated with the skull for 6 weeks, an aluminum halo was mounted on four pins seated in each plate. A titanium can with a plexiglass top to house the Neurochip3 was attached to the halo during all Neurochip3 recording sessions.

Monkey J additionally received EOG implants on the lateral wall and superior margin of both orbits. EOG electrodes consisted of a titanium washer ((#4, 0.25” OD, 0.032” thick) with a 0.016” hole drilled into it and a 34-gauge silver-plated copper microwire with silicone shielding (Cooner Wire, AS155-34) threaded through the hole and soldered to the washer. An incision was made to the dorsal and lateral margins of both orbits to expose the skull. A small hole was drilled in each incision, a titanium skull screw (2 mm x 6 mm) was used to hold down the EOG electrodes, and the wires of the electrodes were tunneled beneath the tissue along the skull. Another incision was made along the top of the skull to secure a pedestal with 8 skull screws and allow access to the electrode wires. The inside of the pedestal was filled with surgical silicone adhesive (Kwik-Sil, World Precision Instruments) to prevent infection. Connectors were fastened to the wires postoperatively.

After each surgery animals received postoperative courses of analgesics and antibiotics. Animals did not show signs of discomfort or pain related to any implanted devices after recovery, and all exposed implants were regularly disinfected biweekly with chlorohexidine and treated with antibiotics to prevent infection.

#### Electrophysiology

All data was collected with the head-mounted Neurochip3 (Shupe et al., 2021) while the monkeys were freely behaving in their home cage (Figure 1A). The Neurochip3 is a battery powered bidirectional brain-computer interface capable of saving data to an SD card, allowing for wireless recording and stimulation for up to 24 hours. 16 channels of the Utah array, or 14 channels of the array and 2 channels of the EOG, were recorded at 20 kHz sampling rate with a bandwidth of 0.1 Hz to 5 kHz (Figure 1B). The cortical channels were chosen each day with a preliminary recording to capture the largest single units present in the array. Most experiments recorded a different set of channels, depending on best spike recordings.

**Figure 1.**
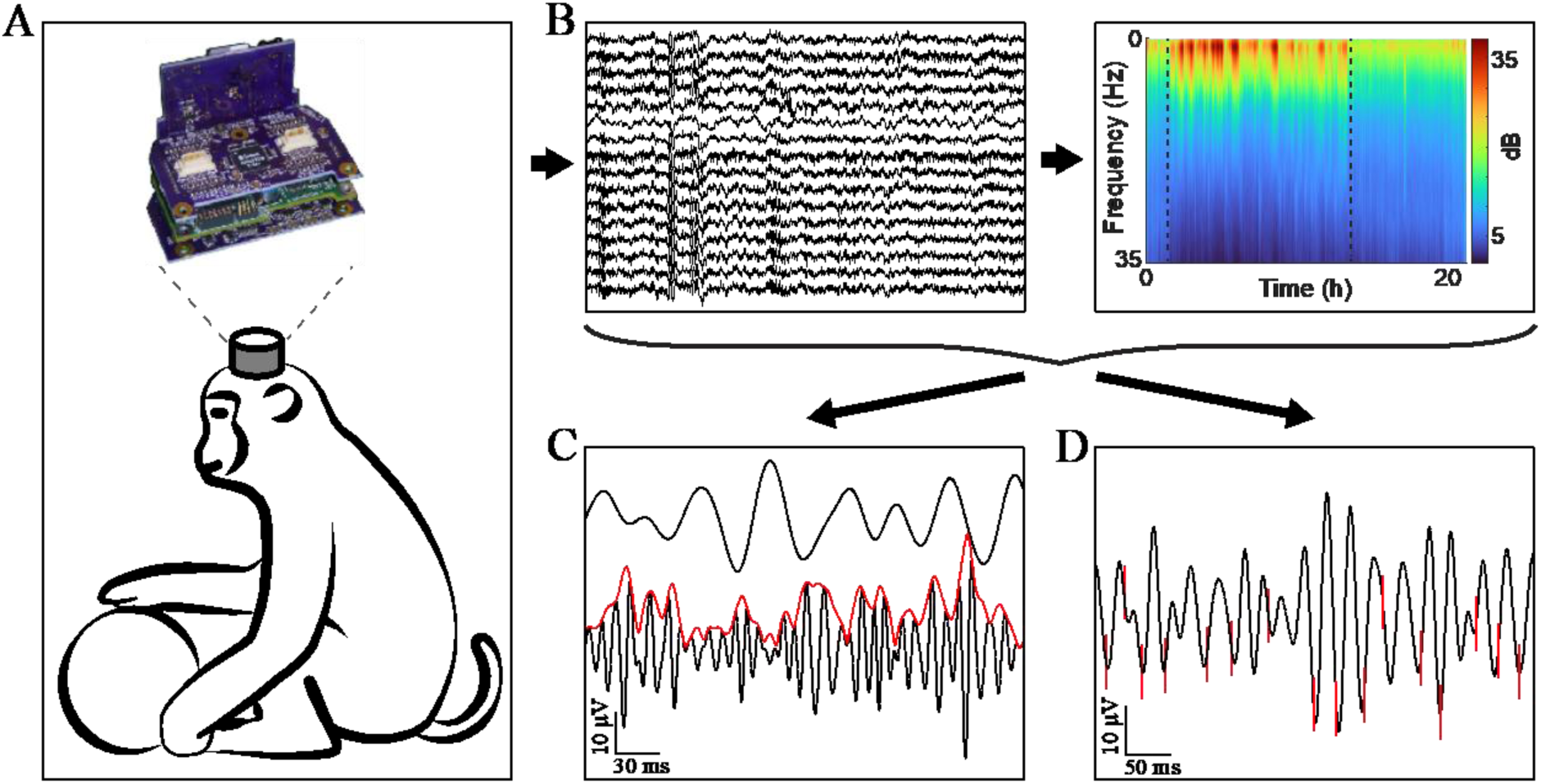
Experimental design. **A.** All data was gathered using the Neurochip3 on freely behaving macaques for up to 24 hours. **B.** We collected 16 channels of data at 20 kHz and obtained the spectral density of every 8-second bin to classify behavioral states. **C.** We then calculated the cross-frequency phase-amplitude coupling of every frequency band pair. The example shows the instantaneous amplitude (red) of high gamma filtered LFP (black, bottom) increasing at the trough of beta (black, top). **D.** We additionally sorted single units and found the synchrony of spikes to LFP bands. The example shows spikes (red) synchronized with the trough of beta filtered LFP.

The Neurochip3 also has an onboard 3-axis accelerometer that was simultaneously recorded at 100 samples per second. Recording sessions lasted between 19 and 24 hours. The lights in the animal rooms were off during the night for the 12 hours between 6pm and 6am. We collected data from a total of 16 sessions from Monkey K and 10 sessions from Monkey J.

### Data analysis

#### Classifying sleep states

All analyses were performed using custom MATLAB (Mathworks) and Python code. The power spectral density (PSD) of cortical local field potentials (LFPs) as well as the onboard accelerometer data from the Neurochip3 were used to classify different sleep states (Figure 1B). Data was down sampled to 1 kHz before performing Welch’s PSD estimate between 0 and 50 Hz for each 8 second time-bin. We then converted the PSD into power and found the average power across all channels for each time-bin. The average power was further normalized by subtracting the minimum value and dividing by the integral to ensure that relative power at different frequencies played a larger role than the absolute power.

We then used the normalized average power as inputs to train a stacked sparse autoencoder for dimensionality reduction, similar to the architecture described in Tsinalis et al. 2016 (Tsinalis et al., 2016). The encoder was composed of 3 layers of 256, 128, and 64 units each, with batch normalization and ReLU activation function. The final hidden layer containing the reduced representation of the data had 32 nodes. The autoencoder was trained with minibatch sizes of 64 for 300 epochs. The loss was calculated with mean squared error with L1 regularization (regularization weight *λ* = 1*e*^−5^) using the Adam optimizer (learning rate *α* = 1*e*^−3^, decay rate for first moment *β*_1_ = 0.9, decay rate for second moment *β*_2_ = 0.999, constant *ε* = 1*e*^−8^). The autoencoder was implemented in Python using the PyTorch package (Paszke et al., 2019).

Accelerometer data was included by performing the root sum of squares across all three axes. We then found the variance of the values within each time bin and applied a logarithmic scale to better compress the data. Finally, the standard deviation was normalized to that of the encoded dimension with the largest variance. The processed accelerometer data was included as an additional dimension in the lower dimensional representation (i.e., as the 33^rd^ dimension).

To classify the data, we used k-means clustering with an assumption of 4 centroids. Data points of the lower dimensional representation within the 90^th^ percentile of pairwise Euclidean distance were initially used for finding the centroids of clusters to avoid the influence of outliers. Each data point was assigned to the centroid with the shortest Euclidean distance.

After clustering, each group of records was assigned to one of four states – 1) awake and moving (Move) 2) awake and at rest (Rest), 3) rapid-eye movement (REM) sleep, 4) non-REM (NREM) sleep – by assessing the average accelerometer value and average normalized PSD for each cluster. First, the cluster with highest average acceleration was assigned to be Move, then the cluster with highest average delta power (0.5 to 4 Hz) was assigned be NREM, then the cluster with higher average beta power (15 to 30 Hz) was assigned to be Rest, and the remaining cluster was assigned to be REM.

To include a temporal aspect and smooth any outliers we performed a majority filter on the classification. For each time-bin, the majority state across 2 time-bins before to 2 time-bins after (±16 seconds) was considered to be the current state. Ties were resolved by keeping the original classification, or by random choice if the original classification was not part of the tie.

#### Validation of classification

Two EOG electrodes (one dorsal and one lateral) were simultaneously recorded from with the Neurochip3 in experiments with Monkey J. EOG signals were extracted by subtracting the lateral electrode signal from the dorsal electrode signal and then applying a band-pass filter between 1 and 20 Hz with a zero-phase second-order Butterworth filter. We performed in-booth experiments with flashes of lights guiding the monkey’s gaze to ensure we were properly capturing eye movements.

As the Neurochip3 is mounted to the monkeys’ heads, we performed overnight recordings with the Microsoft Kinect in conjunction to ensure there were no large discrepancies between the on-board accelerometer and whole-body movements (Libey & Fetz, 2017). Movements with the Kinect was calculated as the absolute difference between each frame of the infrared depth-finding camera and the previous frame. We captured frames as quickly as possible with the processing overhead, around 30 frames per second.

Further validation was performed with k-fold cross-validation with k=20. Each dataset was split into 20 random groups; classification was performed on 19 of the 20 groups, and the final group was classified using the centroids from the classification k-means. The training error was calculated as the difference in classification within the 19 groups used for training the classification, and the test error as the difference in classification within the final group used to test the classification. Cross-validation was bootstrapped 50 times for each session to minimize variability that can potentially be introduced by the random sampling.

#### Coherence

Magnitude-squared coherence was used to calculate synchrony within the same frequencies:

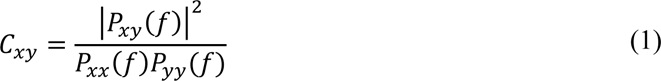

where *C_xy_* is the coherence between *x* and *y*, *P*_*xy*_(*f*) is the cross-spectral density between *x* and *y*, and *P*_*xx*_(*f*) and *P*_*yy*_(*f*) are the spectral densities of *x* and *y* respectively. Coherence was calculated between all combinations of channel pairs at every 0.1 Hz intervals.

#### Cross-frequency phase-amplitude coupling

To calculate cross-frequency phase-amplitude coupling we used mean vector length (MVL) (Canolty et al., 2006) (Figure 1C). The LFP was first filtered into 2 frequency bands of interest within the 6 frequency bands of interest – 1) delta (0.5-4 Hz), 2) theta (4-8 Hz), 3) alpha (8-12 Hz), 4) beta (15-30 Hz), 5) low gamma (30-70 Hz), and 5) high gamma (70-120 Hz) – using a zero-phase second-order Butterworth filter. We then calculated the analytic signal, *H*, of each band using the Hilbert transform:

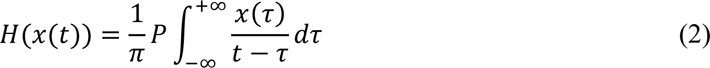

where *P* is Cauchy principal value. The phase of the complex valued analytic signal is the instantaneous phase at time *t*, and the magnitude of the analytic signal is the instantaneous amplitude. The mean vector was then calculated by:

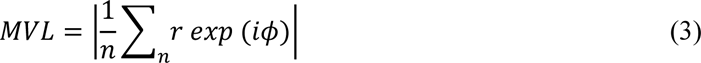

where *r* is the instantaneous magnitude of the higher frequency band and *ϕ* is the instantaneous phase of the lower frequency band. *ϕ* ranges from 0 to 2*π* radians where 0 is the peak and *π* the trough of oscillations. This calculation creates a vector for each sample in time with the phase and amplitude of the lower and higher frequency bands, respectively. The magnitude of the average of those vectors, or MVL, measures the strength of synchrony – zero indicates a uniform distribution in which the vectors “cancel” each other, and higher values indicate the degree of synchrony.

However, MVL is highly affected by the amplitude and does not have a normalized maximum value, which makes interpretation of individual values and comparisons of MVL measurements across different time points difficult. Thus, we additionally calculated the maximum possible MVL for each state. Instead of using the phase and amplitude that occurs at the same time sample in Equation 2, we paired the highest amplitudes with the most commonly occurring phases. Thus, the largest vectors were all in similar directions providing the maximum possible length of the mean vector. We then normalized the MVL by dividing by the maximum possible MVL to assess differences in synchrony between behavioral states.

#### Spike sorting

Spikes were sorted offline using two-window discrimination. The cortical recording was bandpass filtered between 1000 and 2000 Hz with a first-order Butterworth filter. Then a negative threshold and two time-delayed windows were manually chosen to capture the trough and the peak of the spike waveform. All traces crossing the threshold and passing through the two windows was denoted as a spike.

As the fidelity of spikes may change over such a long period of recording, we additionally compared the shapes of the first 1000 detected spikes with the last 1000 detected spikes using the coefficient of determination (CoD):

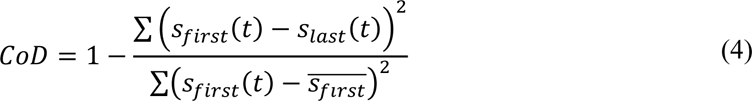

where *s*_*first*_ is a waveform of one of the first spikes, *s*_*last*_ is a waveform of one of the last spikes, and 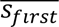 is the average of the waveform.

We performed pairwise CoD on all first and last 1000 instances of each spike and compared them to the pairwise CoD between the first 1000 instances of the spike and the last 1000 instances of a different, randomly chosen spike. If the distribution of the CoD comparing the same spike was significantly higher than the distribution between the spike and another spike it was considered to be consistent overnight. Additionally, to ensure we were not capturing multiple neurons with a similar waveform, we manually assessed the autocorrelograms for each spike to ensure the presence of a refractory period and proper distribution of inter-spike intervals.

#### Phase-locking value

We calculated the phase-locking value (PLV) to assess the strength of synchronization of spike timing to phases of oscillations in specific frequency bands from LFPs recorded on the same channel (Figure 1D). First, the LFP was filtered into a frequency band of interest. The instantaneous phase of the lower frequency band was calculated as described above. We then calculated the PLV:

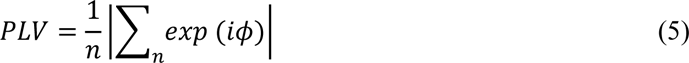

where *ɸ* is the instantaneous phase at spike times. The PLV effectively converts each phase into a unitary vector and finds the average of all the vectors. The magnitude of the resulting vector determines the synchrony of the phases. A value of zero, similar to MVL, indicates a uniform distribution, or no synchrony. A value of one indicates that all phases are equal, or perfect synchrony.

The average phase of spike timing was found using the circular mean:

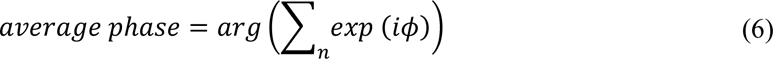

where *arg* is the argument of a complex number, or the angle between the positive real axis and the complex number in the complex plane.

#### Statistical analysis

We used Friedman’s test to compare values between the four states due to the non-parametric nature of the distribution and because we were sampling the same spikes and LFPs across each state. Tukey’s honest significance test was used post hoc to determine significant pairwise differences.

## Results

### Classification of sleep states

Although there have been various approaches automating sleep state classification by taking advantage of the sequential nature of the states as suggested by the American Academy of Sleep Medicine (AASM) classification (Craik et al., 2019; Silber et al., 2007; Supratak et al., 2017), studies have shown that including information from neighboring epochs does not necessarily improve classification (Sekkal et al., 2022; Tsinalis et al., 2016). Most deep learning methods also rely on supervised learning, but, due to the high inter-scorer variability in manual classification that these models rely on (Himanen & Hasan, 2000; Younes et al., 2016), we chose an unsupervised method instead. As a result, we used dimensionality reduction for feature extraction and subsequent clustering. Autoencoders were chosen as the method for dimensionality reduction due to their nonlinearity potentially extracting more salient features compared to linear methods. Denoising autoencoders are often used for feature extraction to ensure the network does not learn to replicate consistent noise, but our input data inherently contained random noise due to the recording device being mounted on a freely behaving animal as well as the short time window of 8 seconds for our power spectral density calculations. As a result, we used a stacked sparse autoencoder for dimensionality reduction.

Sleep states were classified using the power spectral density of local field potentials (LFPs) of successive 8-second time-bins as the input to train an autoencoder. We then extracted the values from the hidden units as the low-dimensional representation of the data. Figure 2A shows examples of the original spectra and reconstructed outputs after training the autoencoder. The fluctuations in the spectra have been smoothed out but the salient features, such as peaks at alpha or beta, were maintained. Figure 2B shows the low-dimensional representation visualized via t-distributed stochastic neighbor embedding (t-SNE) (van der Maaten & Hinton, 2008) after full classification. There was some overlap between the states in the representation, which is to be expected as brain states during sleep are on a continuous spectrum. Finally, we wanted to gain insight into which feature each of the hidden units of the autoencoder represent. Figure 2C shows the average of 100 spectra that gave the largest values in each hidden unit and Figure 2D shows the average of 100 spectra that gave the smallest values in each hidden unit subtracted from spectra in Figure 2C. This analysis can be interpreted as the features within the spectra that each hidden unit encodes, and clear peaks in each frequency band can be observed.

**Figure 2.**
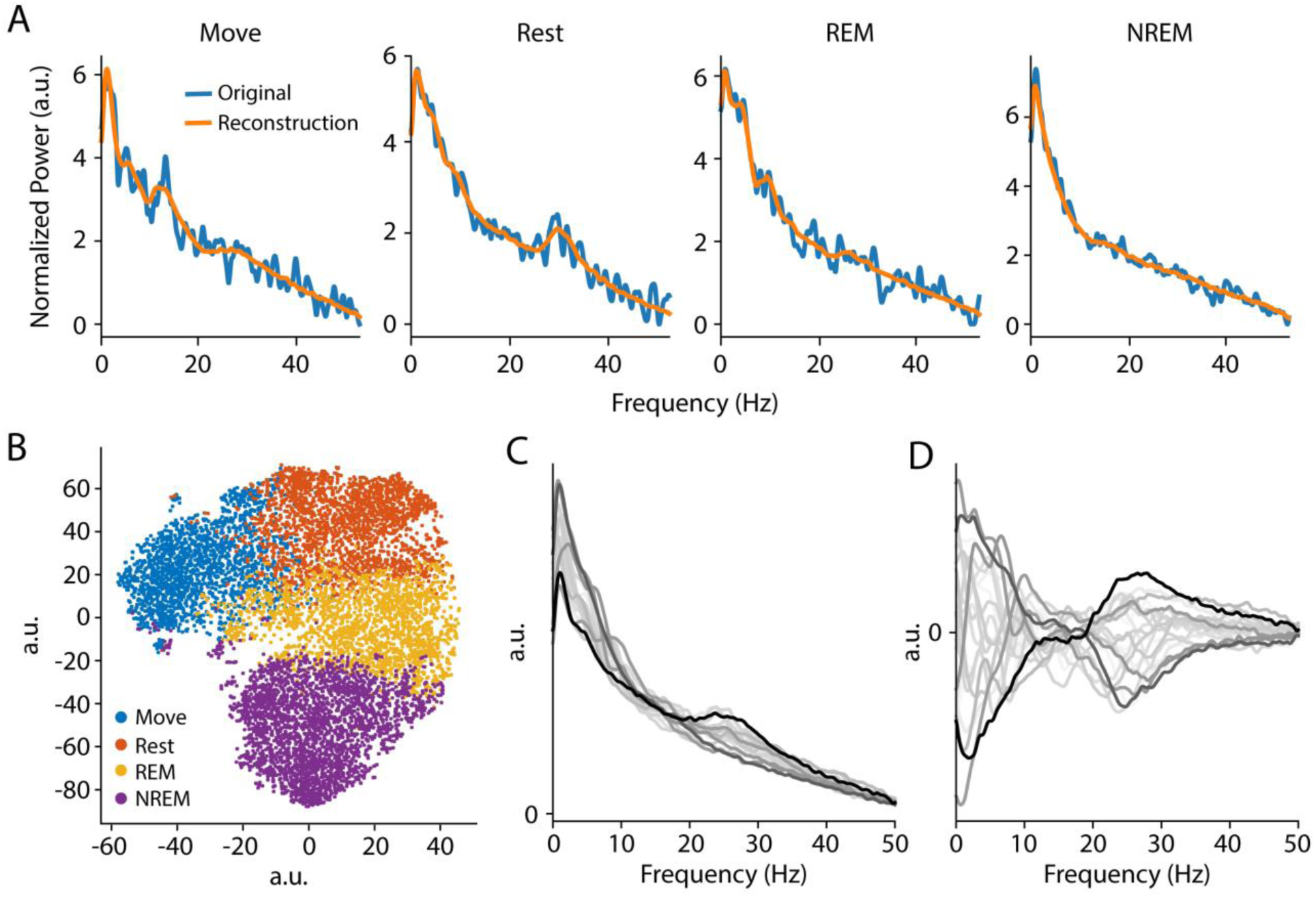
Dimensionality reduction. **A.** Examples of original spectra and the reconstruction at the output of a trained autoencoder for each classified state. **B.** t-SNE visualization of the low dimensional representation with finalized states. **C.** An example of the average of 100 spectra that generated the highest values in each hidden unit. The darkness of the traces is weighted by the variance in the values of all values in that hidden unit (i.e. more relevant during clustering). **D.** The average of 100 spectra that generated the lowest values in each hidden unit subtracted from (C).

We then applied k-means clustering on the lower dimensional representation as well as the accelerometer data that was collected concurrently through the Neurochip3. We assumed 4 clusters – awake and moving (Move), awake and at rest (Rest), REM sleep (REM) and non-REM sleep (NREM). Each cluster was assigned to the state depending on the average accelerometer value and features in the average power spectral density (PSD). Although we have a physiological basis as to why we chose 4 clusters, we wanted to ensure that 4 clusters was a reasonable number for our dataset. To that end, we tracked the within-clusters sum-of-squares as well as the average silhouette coefficient for 1 to 10 clusters (Figure 3A). Both values decrease with more clusters, but the “elbow” point, or where there is a large reduction in the decrease, was around 4 or 5 clusters. We additionally calculated the individual silhouette coefficients when using 4 clusters to show that each cluster had a significant number of data points with coefficients above the average value (Figure 3B).

**Figure 3.**
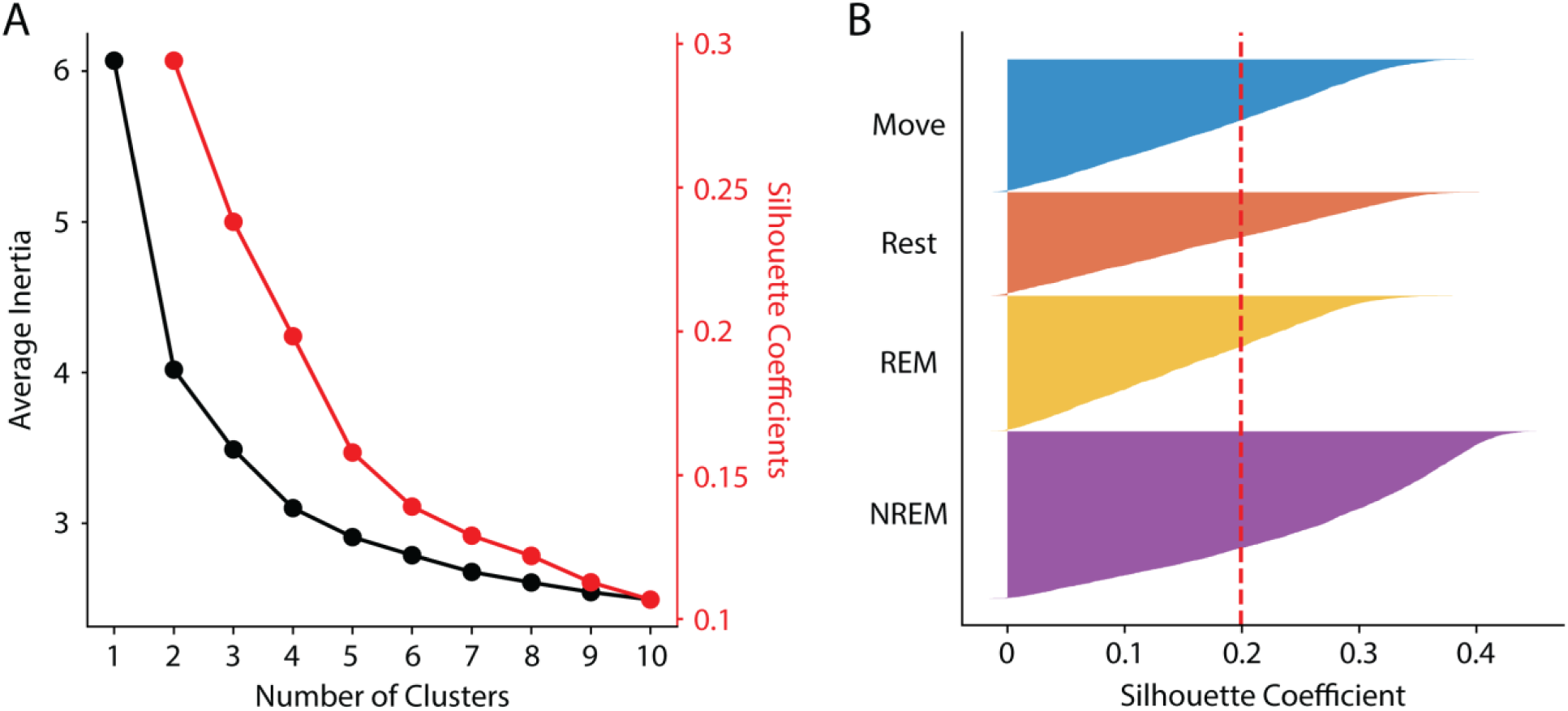
K-means clustering. **A.** The average inertia (black), or within-clusters sum-of-squares, and the average silhouette coefficient (red) of the low dimensional data for different number of clusters. **B.** The silhouette coefficient of each data point with the final classifications compared to the average (vertical red line). All clusters have significant portions above the average.

### Classified states show distinct features

The full classification procedure is outline in Figure 4A and an example of the final classification over a 21-hour period and the corresponding spectral power is shown in Figure 4B. During lights-off we found consistent REM cycles in between NREM epochs occurring every 30 minutes to 2 hours, which became longer and more frequent closer to the morning, consistent with previous findings (Aserinsky & Kleitman, 1953; Hsieh et al., 2008; Kripke et al., 1968). NREM was absent when the lights were on in the animal room, but we often found brief periods of REM sleep concurrent with changes in the PSD, which we attributed to naps (Figure 4B, arrow) (Dijk et al., 1987; Moses et al., 1975). We also often found short periods of Rest or sometimes even Move during the night which were attributed to brief awakenings, typical for a normal night of sleep.

**Figure 4.**
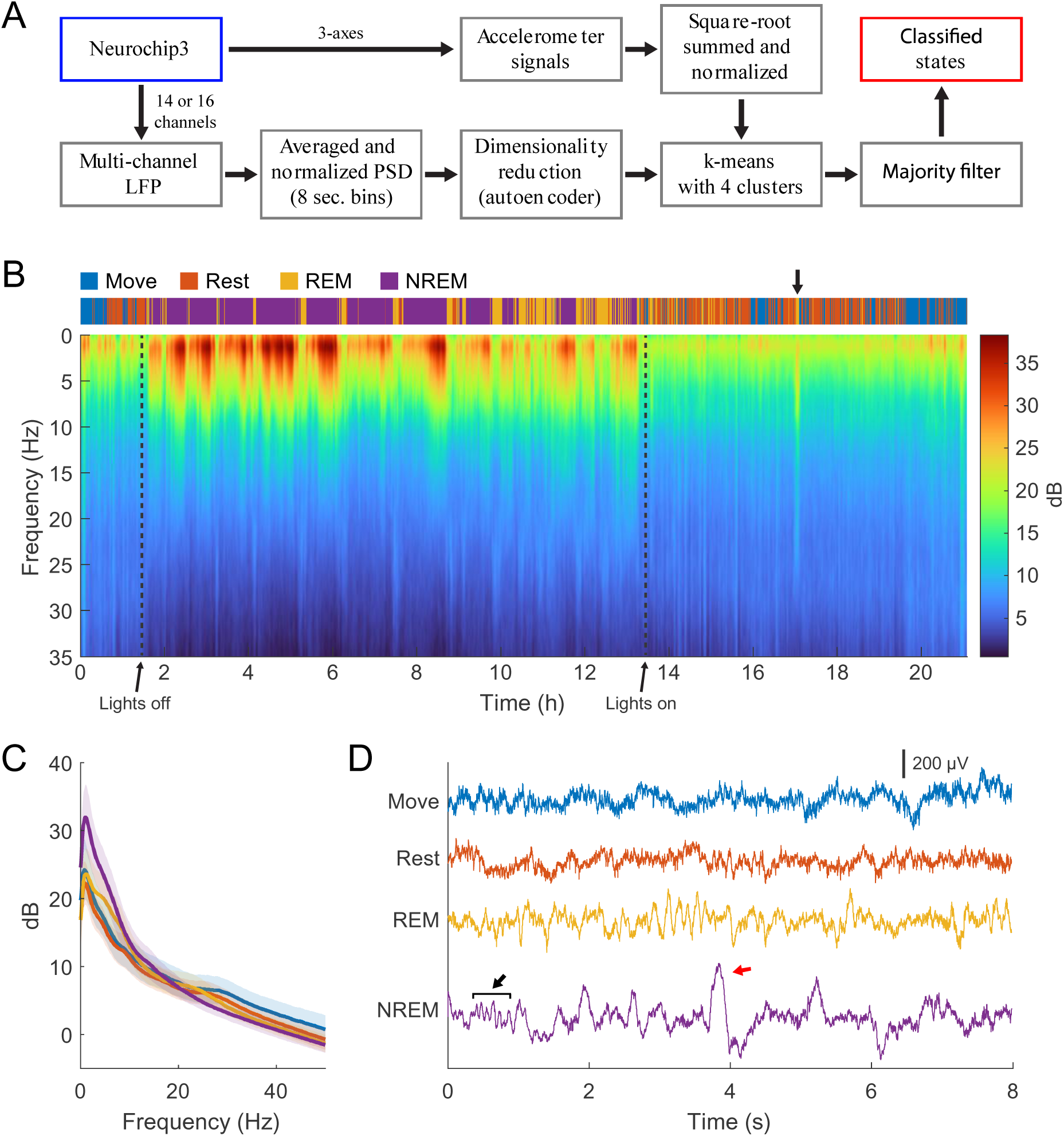
State classification. **A.** Diagram of the full classification process. Normalized LFP power spectral density was used to train an autoencoder for dimensionality reduction. The lower dimensional representation and normalized accelerometer signals were subsequently clustered with k-means clustering then smoothed with a majority filter to determine different sleep states. **B.** Example of classification and spectra over 22 hours of continuous recording. Brief periods of REM sleep were observed during the day, attributed to naps (arrow). **C.** Averaged spectra across each state for the session shown in (B). There is high delta power during NREM, theta power during REM, and beta power during Move and Rest states. **D.** Example traces of raw LFP during each state. Spindles (black arrow) and k-complexes (red arrow) were observed during NREM sleep.

Averaging the spectral power during each sleep state shows high delta power during NREM sleep and high beta power when the animal was awake, as we used these criteria to assign the classified clusters to the appropriate states, but also shows high theta power during REM sleep which was not explicitly a part of the classification scheme (Figure 4C). Additionally, raw LFP traces during the states show high-frequency activity during Move, faster oscillatory activity during Rest in the beta range, slower oscillatory activity during REM sleep in the theta and alpha range, and clear K-complexes and sleep spindles along with low frequency activity around delta during NREM sleep (Figure 4D). T

### Validating sleep state classification

As our classification scheme was unsupervised, we simultaneously recorded electrooculography (EOG) signals to validate the identified sleep states (Figure 5A). Eye movements detected with the EOG were clearly associated with identified sleep states; EOG variance was very high when the animal was awake, very low during NREM sleep, and elevated during REM sleep (Figure 5B). This relationship was present even during very brief windows of detected awakening during the night and naps during the day, indicating high accuracy of state classification.

**Figure 5.**
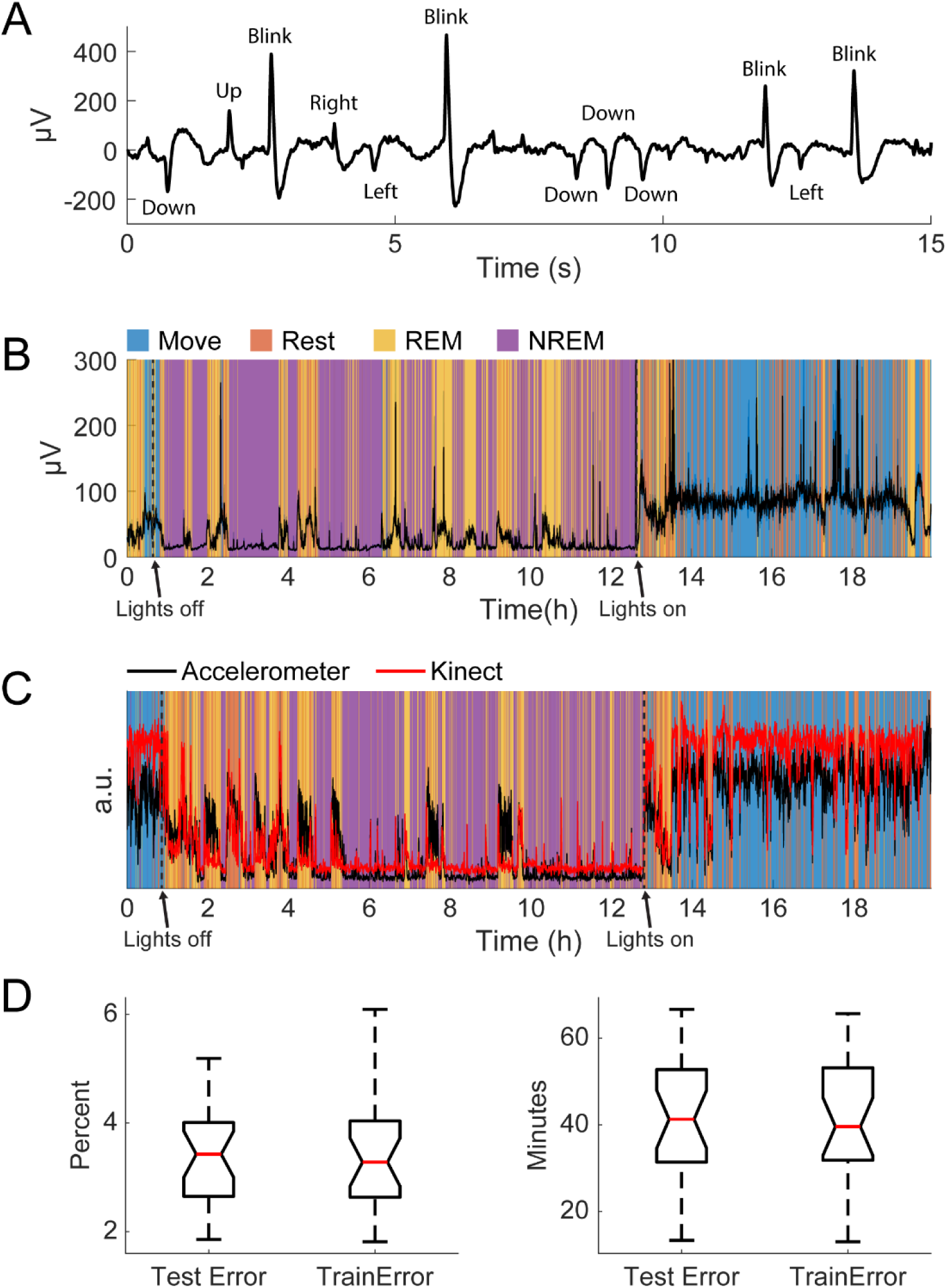
Validation of classification. **A.** An example of filtered differential EOG signals and corresponding eye movements. **B.** Standard deviation of EOG signals overnight with respect to classified states. Note the increase in large eye movements during REM and waking. **C.** An example of the normalized variance of accelerometer data and the normalized variance of the Kinect data with respect to classified states. **D.** Test and training errors with k-fold cross-validation in percent (left) and total minutes (right).

The accelerometer used for classification was on the Neurochip3 which was mounted on the animals’ head. To confirm the accuracy of detected movements, we additionally recorded the monkeys’ movements overnight with a Microsoft Kinect. Compared to the movement values extracted with the Kinect, the accelerometer values were smaller when the animal was awake but larger when the animal was asleep (Figure 5C). This is likely due to body movements that are independent of the head (i.e., isolated limb movements) and are not as accurately recorded in the accelerometer. On the other hand, when the animal is laying down at night most movements involve the head. However, these differences were minor, and the two signals were comparable throughout the recording duration.

Finally, we also performed k-fold cross-validation to verify that our classification method was consistent and robust. We used k=20 and carried out 50 repetitions per session and found the average test and training error to be less than 4%, or around 40 total minutes (Figure 5D). The similar but low test and training errors suggest the classification method had low variance (i.e. not overfit) and high consistency. As a result, the classification was deemed appropriate for the aims of this study.

### Changes in LFP dynamics

The average spectral power per state for each animal is shown in Figure 6A. Delta power was high during NREM sleep due to the presence of slow waves. There was a beta peak during the wake states, though the specific frequency range differed between the animals: 25-30 Hz for Monkey K and 15-20 Hz for Monkey J. REM sleep showed slightly higher theta and alpha power compared to the wake states.

**Figure 6.**
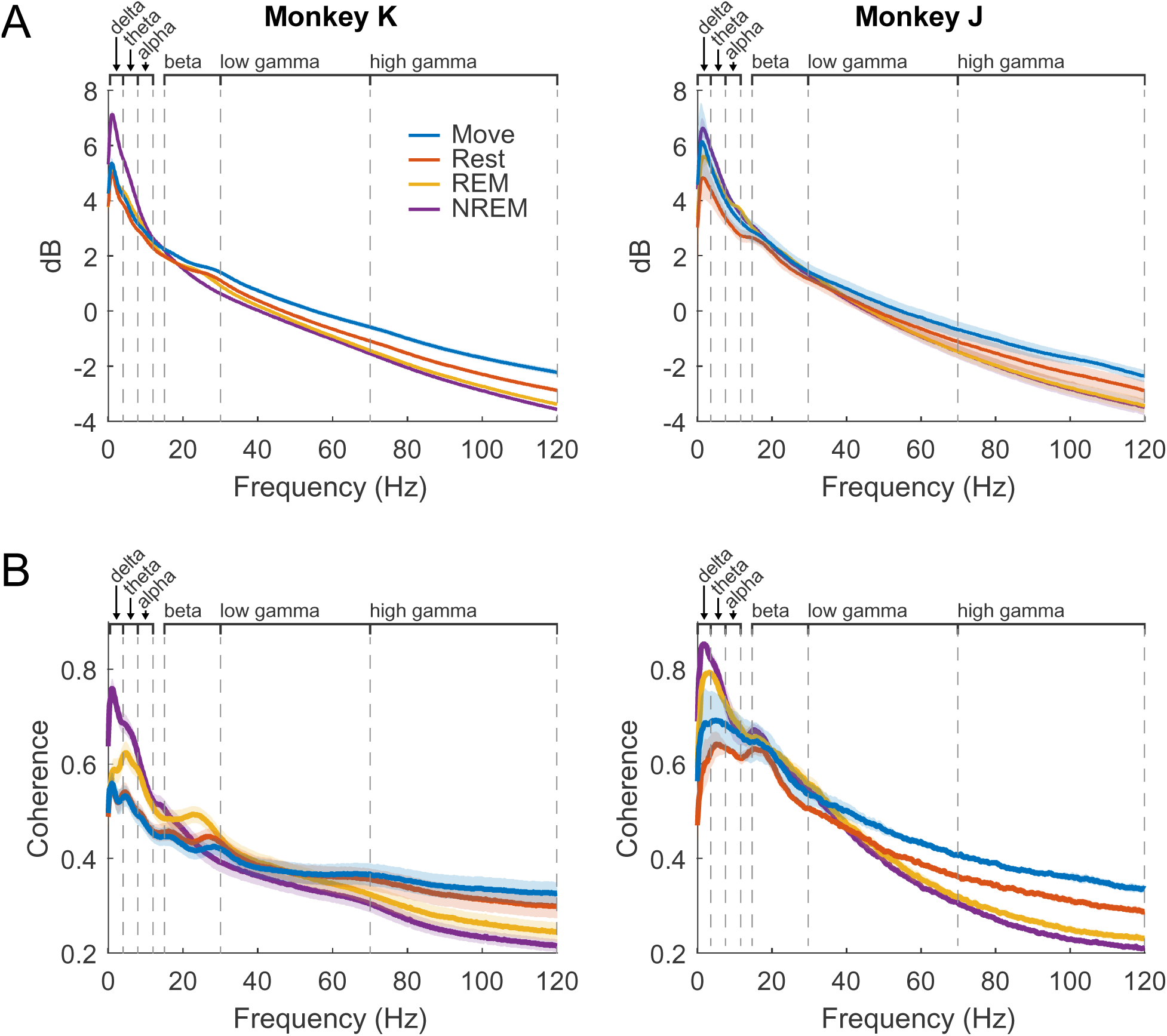
LFP dynamics. **A.** Average spectral power in each state for each animal across all experiments. Shaded regions show standard error. **B.** Average pairwise coherence in each state for each animal across all experiments. Shaded regions show standard error.

Average pairwise coherence per state for each animal is shown in Figure 6B. The features were very similar to those show in the power, but the differences were magnified, likely due to coherence showing synchrony between pairs of channels and amplifying active oscillatory signaling over baseline spectral density. In addition to the large delta peak during NREM, there was a much clearer difference between REM and the wake states, including a more distinct beta peak.

### Cross-frequency phase-amplitude coupling

The phase of lower frequency bands has been observed to be coupled with the amplitude of higher frequency bands, thought to reflect coordination between brain networks (Canolty & Knight, 2010; Jensen & Colgin, 2007). We explored the coupling between every pair of low frequency band phase to high frequency band amplitude to determine whether there were state-dependent changes. We first plotted the average z-scored spectral power of higher frequencies at different phases of each frequency band (Figure 7). There was clear coupling between different phases and frequencies, the strength and specific phase of which changed, depending on the state.

**Figure 7.**
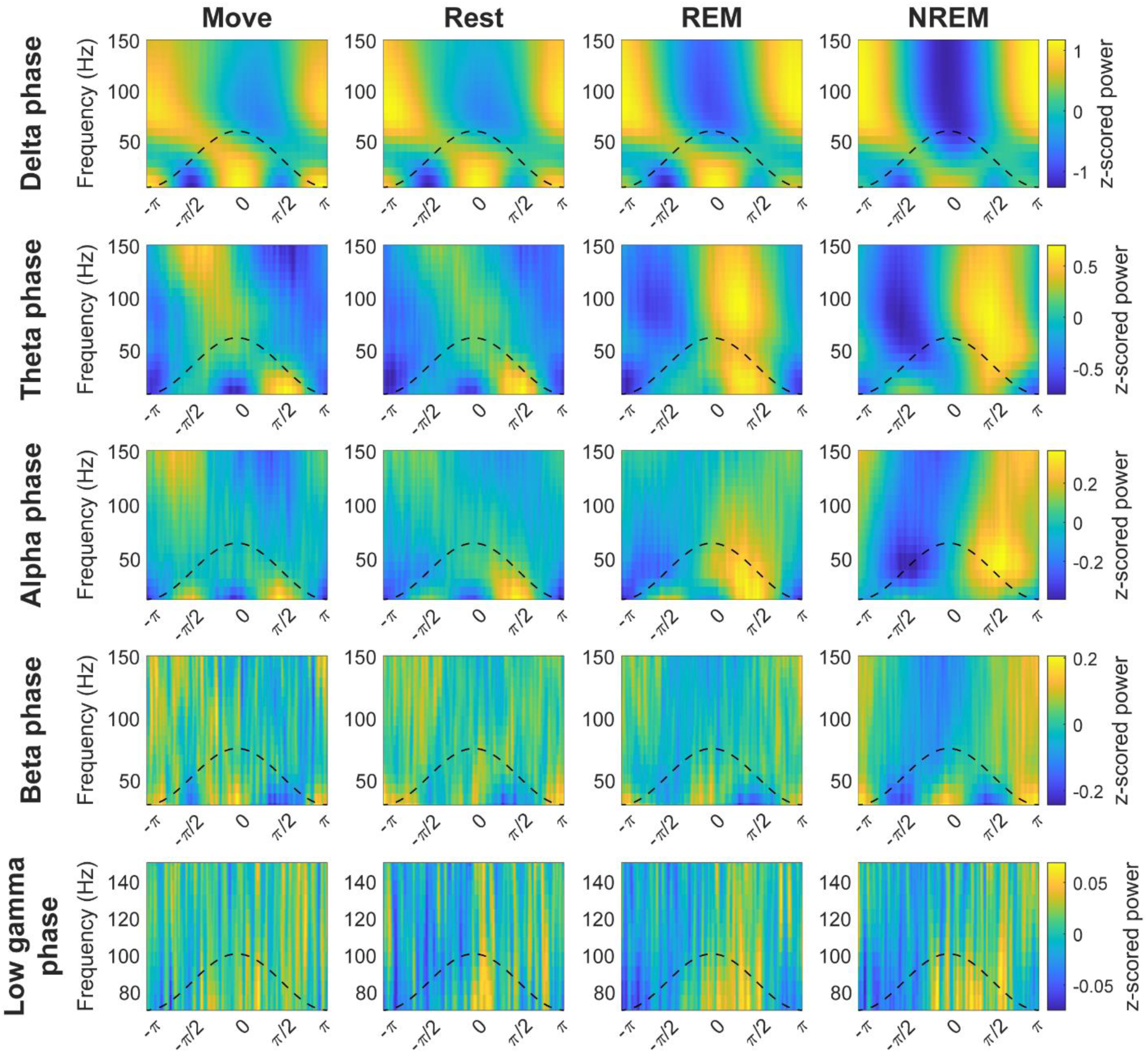
Cross-frequency phase-power distributions. Distribution of lower frequency band phase and higher frequency band power during each state. The power is z-scored for each frequency.

To quantify the degree of coupling we calculated the normalized mean vector length (nMVL). Figure 8 shows the distribution of nMVL for each pair of frequency bands during each state. The distribution of nMVL did have state-dependent changes, notably 1) high delta phase to alpha amplitude (delta-alpha) coupling during Rest, likely due to increased alpha oscillatory power, 2) low delta-beta coupling during NREM, perhaps for disinhibition of the cortex leading to increased plasticity, 3) high delta-high gamma coupling during Move and NREM, potentially reflecting synchrony of spike activity with delta oscillations, 4) high theta-beta coupling during REM, showing a possible modulation of beta by deeper brain structures, 5) high theta-high gamma coupling during Move, which may reflect hippocampal place cells synchronizing cortical activity during behavior, and 6) high alpha-high gamma coupling during Move, which may be due to the relevance of alpha to both attention and movement preparation.

**Figure 8.**
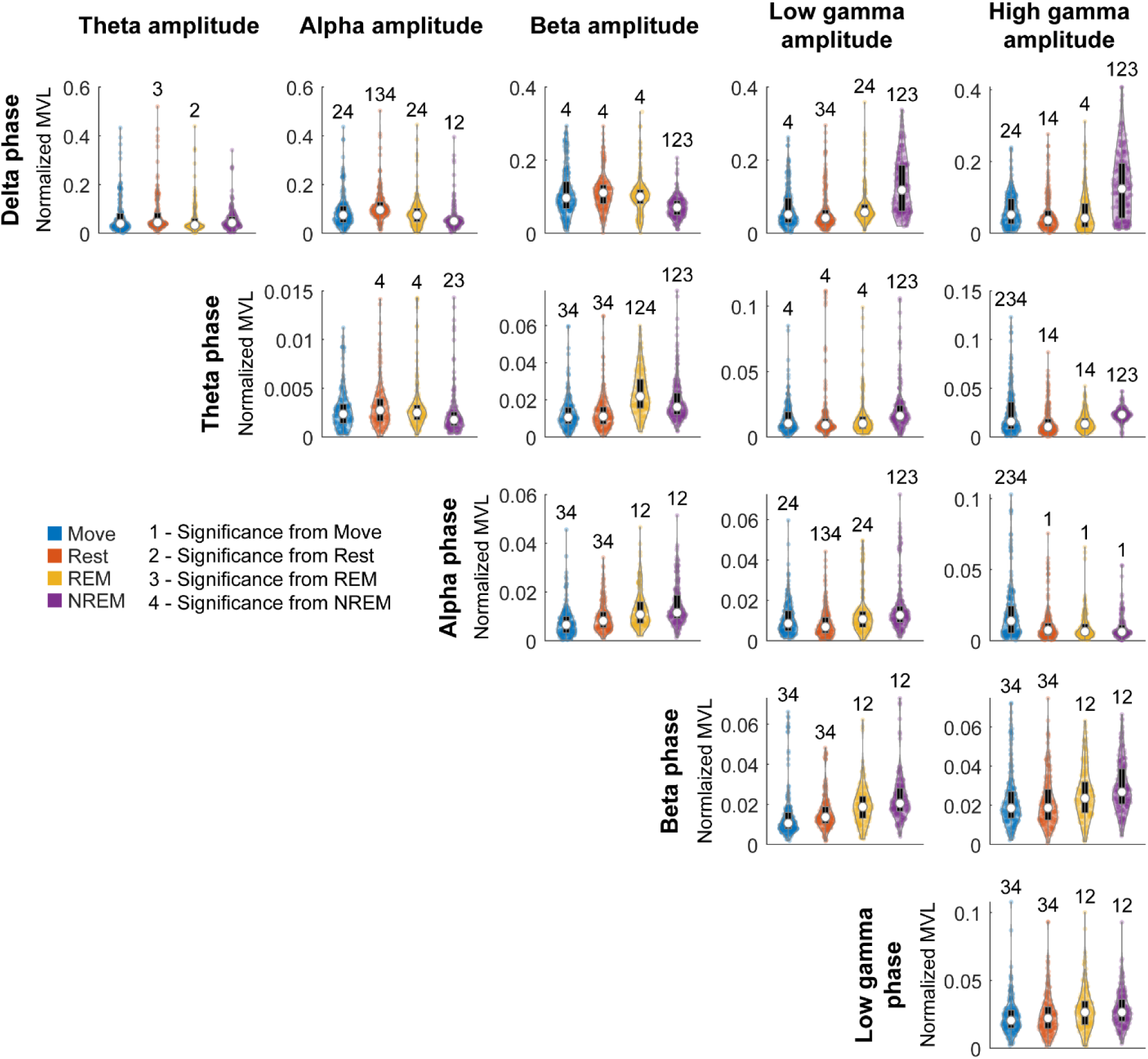
Normalized MVL distributions. Normalized MVL distributions for each lower frequency band phase and higher frequency band amplitude pair during each state. The black boxes show standard box plots with interquartile range and the white dots show median values. The numbers above each group denote significance compared to another state (Friedman’s test, p<0.05).

To further characterize the coupling, we explored the distribution of the mean vectors to determine whether the phases aligned with the highest amplitudes were consistent. Figure 9 shows the distribution of mean vector phases for each pair of frequency bands during each state. The mean phases are consistent across most frequency pairs across all states with stronger consistency typically in aforementioned significant nMVL values (e.g. delta-high gamma during NREM or theta-beta during REM).

**Figure 9.**
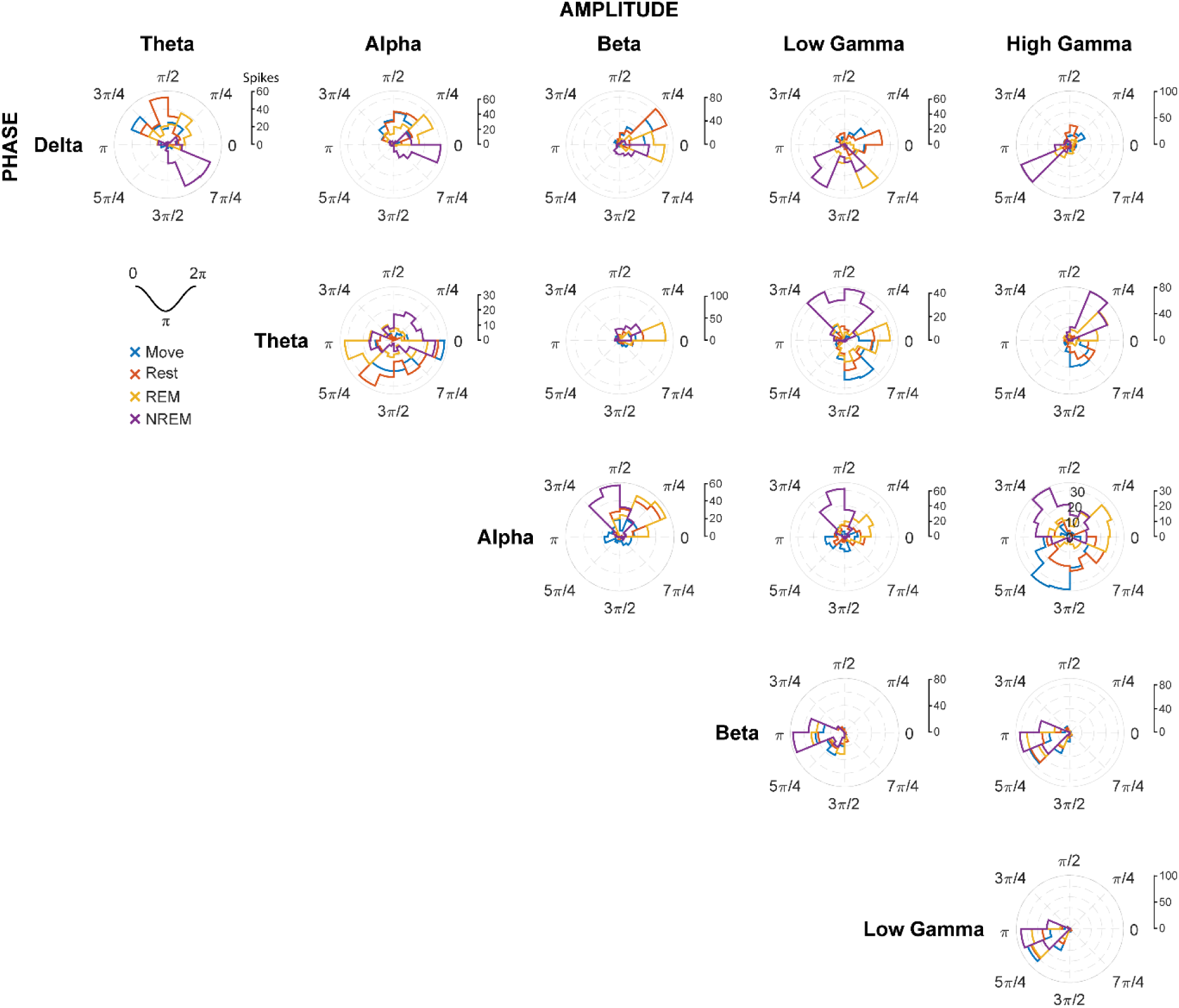
Mean vector phase distributions. The distribution of phases of mean vectors for each lower frequency band phase and higher frequency band amplitude pair during each state. The phase of the mean vector shows the phase at which the amplitude of the higher frequency band is the greatest.

### Spike sorting

Spikes were manually sorted using two-window discrimination and confirmed using the coefficient of determination (Figures 10A and 10B, see *Materials and Methods – Spike sorting*). We tracked a total of 193 spikes (121 in Monkey K, 92 in Monkey J) that satisfied our conditions across all sessions. The firing rates of spikes overnight were strongly and consistently modulated by different identified sleep states, especially during REM cycles during the night, further demonstrating the accuracy of the classification method (Figure 10C). Firing rates were consistently lower with deeper sleep.

**Figure 10.**
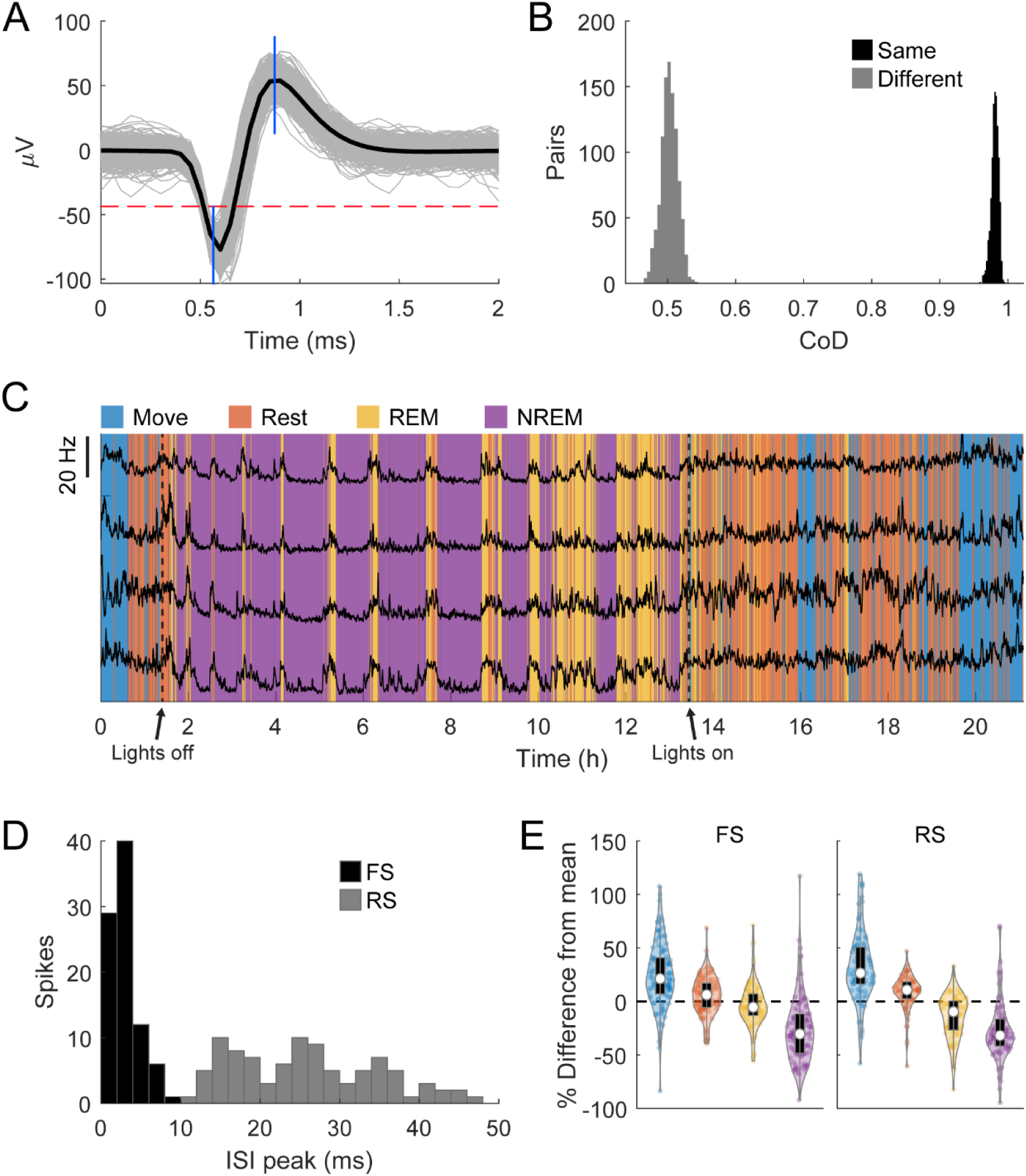
Spike sorting. **A.** An example of a sorted spike. The grey traces show a random sample of 1000 spikes, the black line is the average, the horizontal dashed red line shows the threshold, and the blue vertical lines show the two windows. **B.** An example of comparisons of the pairwise coefficient of determination (CoD) between the first and last 1000 instances of the same spike and the first and last 1000 instances of two different spikes. **C.** Firing rates binned every 60 seconds of four neurons overnight with classified sleep states. The changes in firing rate very closely match the changes in state. **D.** Histogram of the peak of the ISI distribution of each spike. Spikes with peaks earlier than 10 ms were denoted to be fast spiking (FS) and all others denoted to be regular spiking (RS). **E.** Percent difference of firing rate in each state from the average overall firing rate for each spike type. Each state is statistically significantly different from each other state (Friedman’s test, p<0.05).

Additionally, spikes were designated to be regular firing or fast firing. Although spike type classification was traditionally performed with the spike width (McCormick et al., 1985), subsequent studies have suggested that spike width, at least when used alone, may not be the most indicative of the spike type (Insel & Barnes, 2015; Jung et al., 1998; Vigneswaran et al., 2011). Assessment of various features of spikes showed that the peak of the inter-spike interval distribution to be the most distinguishable feature within our dataset (Figure 11). If the peak occurred before 10 ms the spike was designated to be fast spiking (FS), otherwise it was designated to be regular spiking (RS) (84 FS spikes, 109 RS spikes). Assessing the differences in firing rates in individual states showed that both RS and FS spike rates significantly decrease with deeper sleep (Figure 10E).

**Figure 11.**
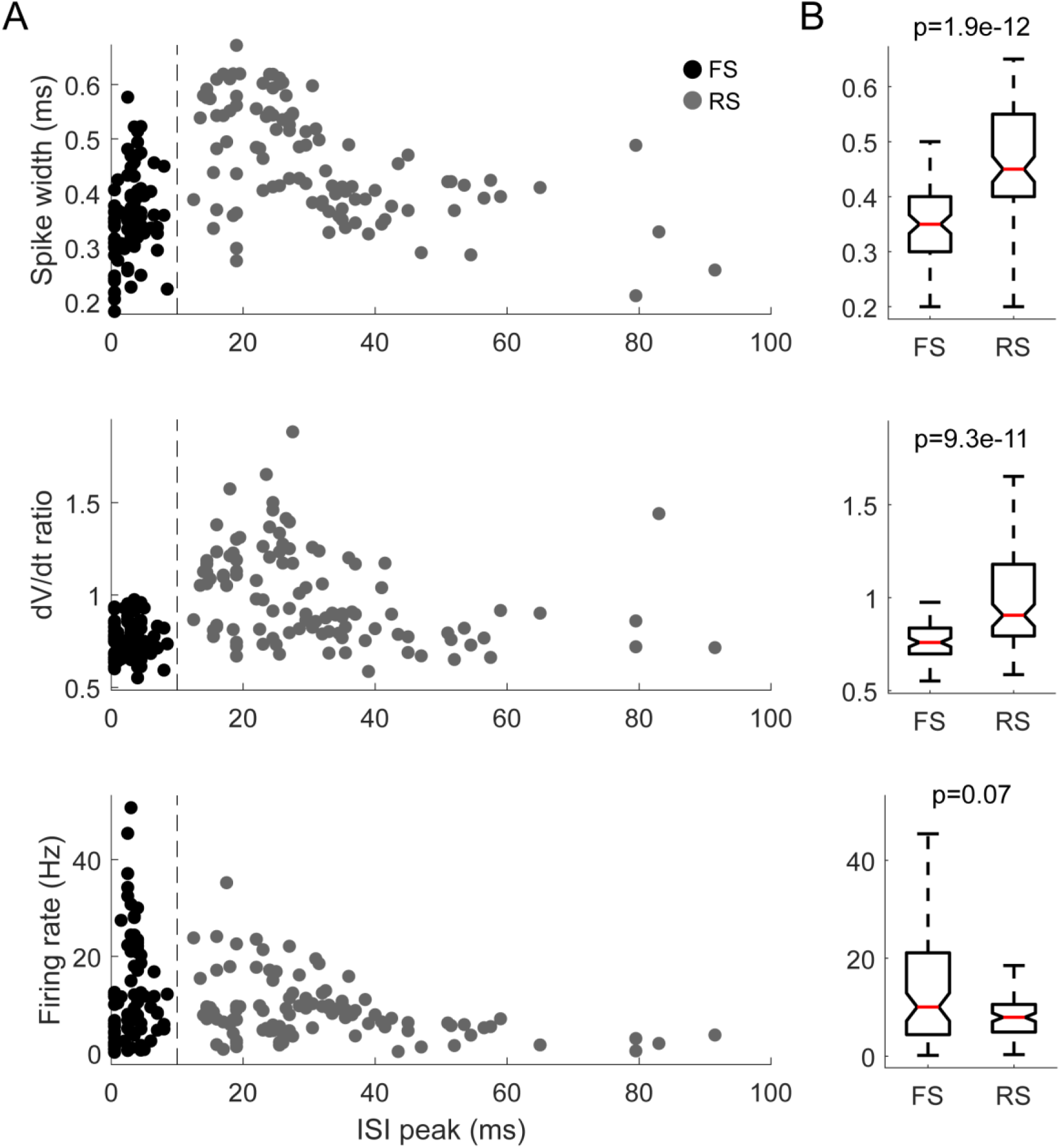
Spike features. **A.** Spike width, dV/dt ratio (the fastest rise of the spike divided by the fastest fall of the spike as defined in McCormick et al. 1985), and the average firing rate with respect to the peak of the inter-spike interval (ISI) distribution. The spike widths include the addition of a random jitter sampled from a uniform distribution from -0.02 to 0.02 ms to better visualize the data. The dashed vertical line denotes the boundary between fast- and regular-spiking neurons (FS and RS respectively) as defined in this study. **B.** Box plots of the corresponding measures and the statistical significance between the two spike types (Wilcoxon rank-sum test). The two measures reflecting the spike shape have significant differences. Although FS neurons have a wider spread of average firing rates, there is no significant difference between the average firing rates of the two types of neurons.

### Changes in spiking dynamics

To determine how spiking patterns changed between different sleep states we analyzed changes in the average ISI distributions of RS and FS spikes. The raw ISI distributions showed an overall decrease on most ISIs with deeper sleep, reflecting the diminished firing rate, but also a slight change in the timing of the distribution, especially for RS spikes (Figure 12A).

**Figure 12.**
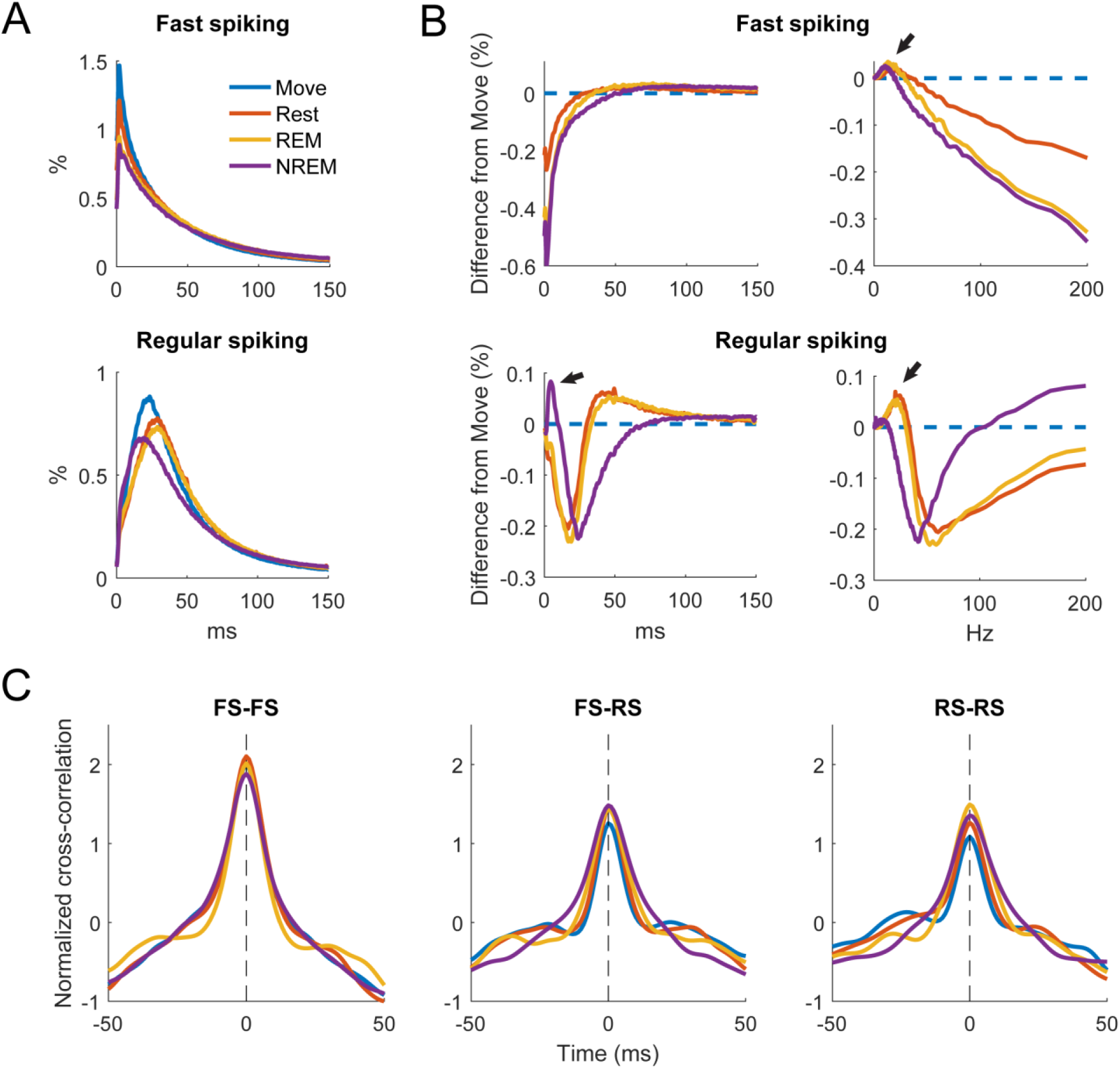
Changes in spiking patterns. **A.** Average ISI distributions of identified fast spiking and regular spiking neurons during each state. **B.** Difference of ISI distributions from Move plotted against time (left) and frequency (right). Note the peak around 5 ms during NREM for regular spiking neurons and the peaks around 7-10 Hz during NREM sleep and the peaks at around 20 and 25 Hz for REM and Rest respectively (arrows) for both neuron types. All three peaks were significantly different from 0 (Wilcoxon signed-rank test, p<0.05). **C.** Cross correlations of spike firing rates between pairs of fast spiking (FS) neurons (left), FS and regular spiking (RS) neurons (middle), and pairs of RS neurons (right) (n = 636, 1328, 1036 respectively).

To quantify the differences, we calculated the difference in ISI distributions from Move (Figure 12B, top). FS spikes had a consistent decrease in shorter latency ISIs with deeper sleep, but RS spikes showed a large increase in short latency ISIs at around 5 ms (arrow). To determine if there were any changes in larger ISIs, we plotted the distributions against frequency (Figure 12B, bottom). Both FS and RS spikes showed a peak between 7 and 10 Hz during NREM and a peak between 20 and 25 Hz during Rest and REM.

We additionally sought to determine how the brain state could affect the relationship between neurons by calculating the cross correlation between firing rates of recorded spikes. Firing rates were calculated by convolving the spike train with a gaussian kernel with an approximate width of 10 ms. Figure 12C shows the cross correlations between smoothed firing rates of pairs of FS neurons, between FS and RS neurons, and between pairs of RS neurons. There was not a clear difference in the cross correlation between the four states suggesting relative circuitry is maintained. The correlation between pairs of FS neurons was stronger than between FS and RS or pairs of RS neurons, as inhibitory neurons are often more interconnected within the cortex.

### Spike-field relationships

Spikes have been shown to be synchronized to various frequency bands, particularly in the motor cortex (Buzsáki et al., 2012; Murthy & Fetz, 1996b). We first analyzed the phase-amplitude distribution normalized by amplitude of each LFP band during spike timings (Figure 13). There was clear synchrony with spikes during NREM sleep with all LFP bands, though at different phases. During NREM spikes typically occurred at the negative phase of delta, beta, and gamma activity, and at intermediate phases for theta (*π*/2) and alpha (3*π*/4). The synchrony was particularly apparent during high amplitudes, suggesting spikes are synchronized to active frequency band activity rather than epiphenomena arising from periodicity of spike firing patterns. During both awake states and REM sleep spikes were synchronized to the trough of beta and gamma frequencies, though less than during NREM sleep. Spikes during Move were also potentially synchronized to the delta band. All other combinations did not show apparent coordination between LFP band phase and spike timing.

**Figure 13.**
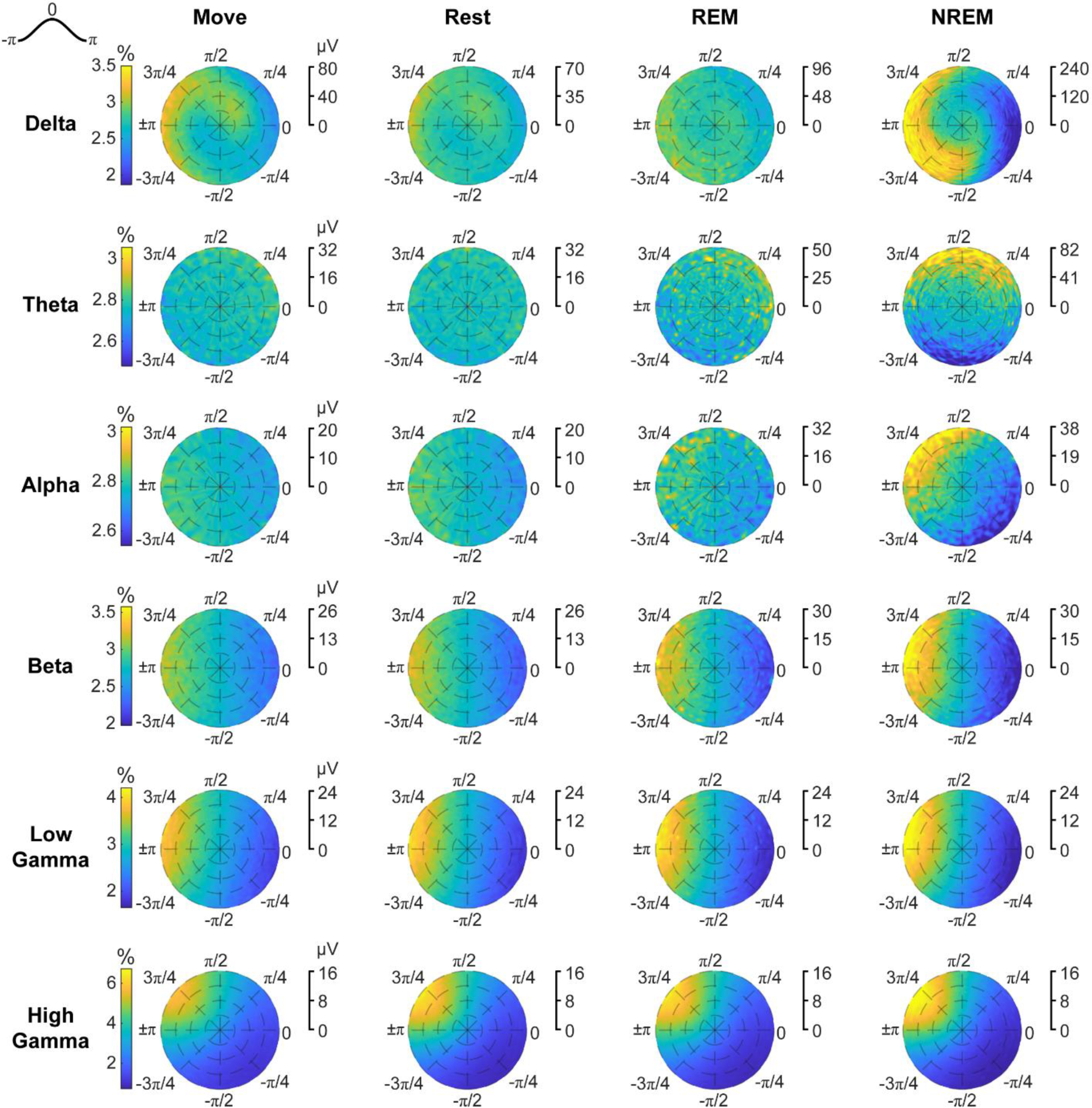
Phase distributions of LFPs at spike times. Phase distributions normalized for each amplitude during each sleep state for each LFP band.

To quantify the strength of these relationships, we calculated the phase-locking value (PLV) for each spike during each state for all LFP bands. Figure 14 shows the RS spikes’ distribution of locked phases as well as the PLV for each frequency band during each state. Spikes were more synchronized to delta during both Move and NREM, validated by the consistency in the locked phases (around *π*/2 and ±*π*, respectively). For every other frequency band, there was stronger synchrony during deeper sleep states, though the differences were often not statistically significant. The phases were extremely consistent for beta and higher frequency bands, but variable for lower frequency bands during states with lower PLVs.

**Figure 14.**
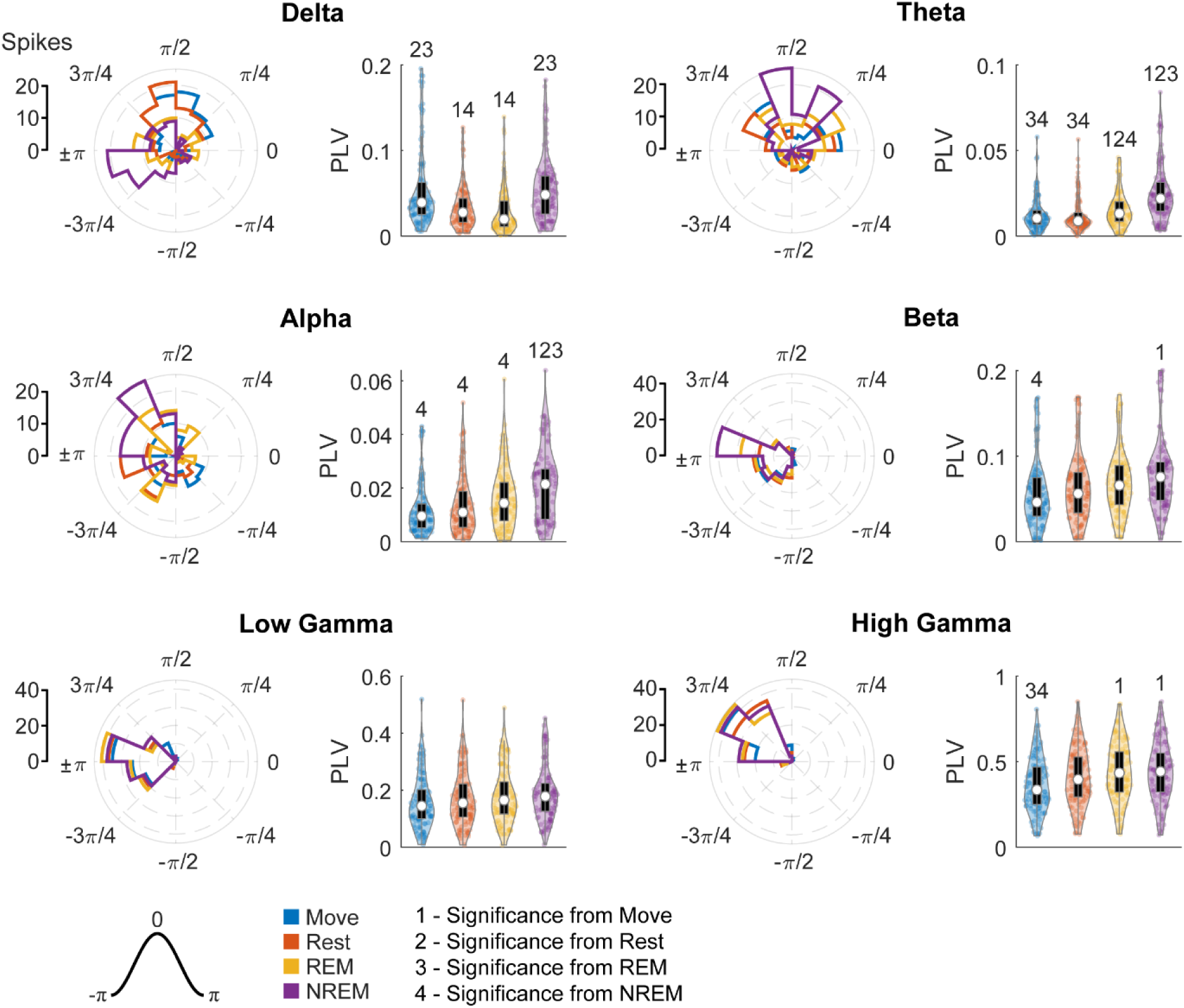
Locked phase and phase locking value distributions for regular spiking neurons. Distributions of locked phase (left) and PLVs (right) of regular spiking neurons during each sleep state for each LFP band. The numbers above each group denote significance compared to another state (Friedman’s test, p<0.05).

Figure 15 shows the FS spikes’ distribution of locked phases and the PLV for each frequency band during each state. One large difference from RS spikes was the synchronization with delta during Move, which was not significantly larger compared to Rest or REM. Otherwise, there was a much stronger relationship between increase in PLV and deeper sleep states, being statistically significant for each frequency band. In addition, the phase of synchrony to the low and high gamma bands was earlier in the phase for FS spikes compared to RS spikes.

**Figure 15.**
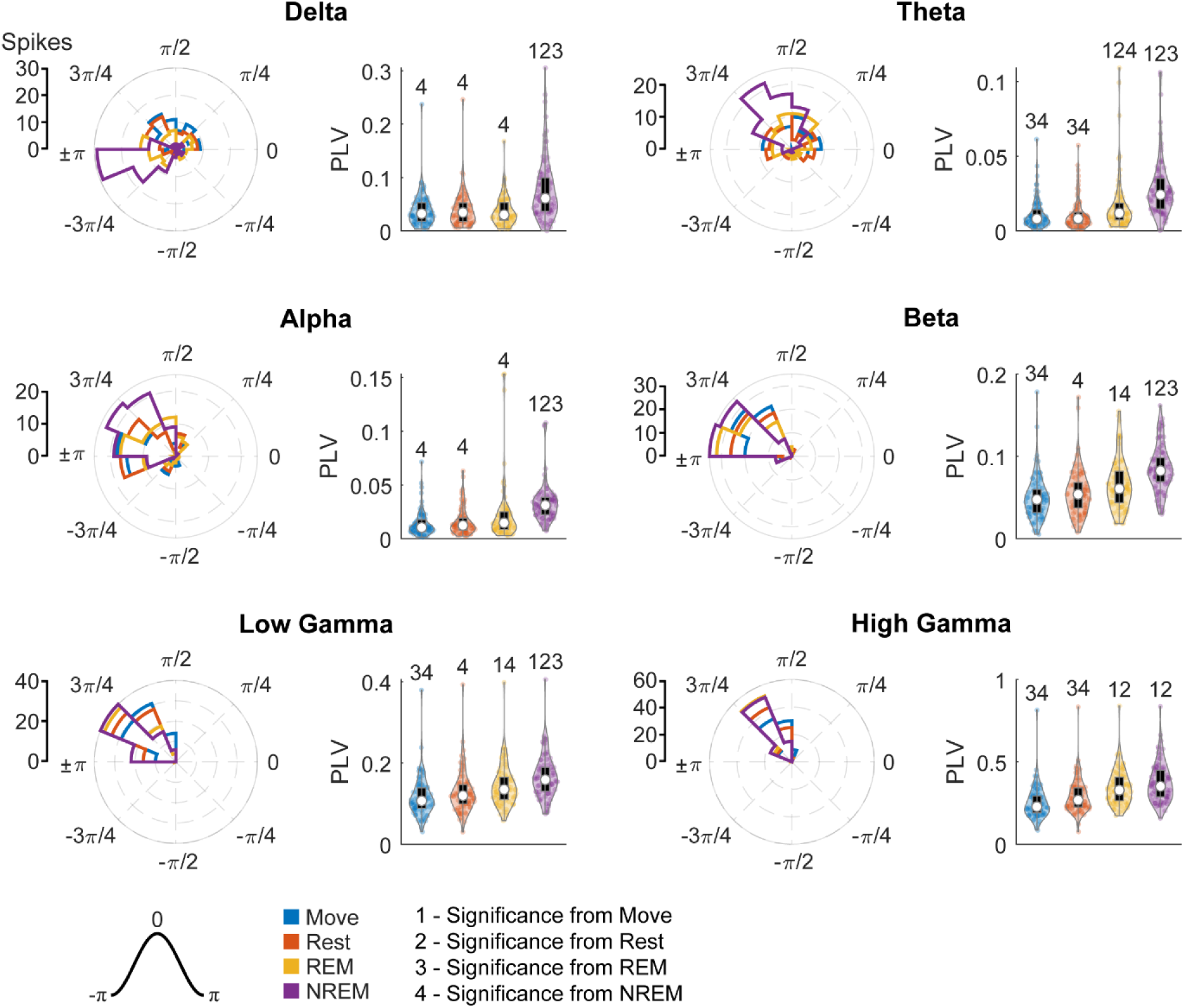
Locked phase and phase locking value distributions for fast spiking neurons. Distributions of locked phase (left) and PLVs (right) of fast spiking neurons during each sleep state for each LFP band. The numbers above each group denote significance compared to another state (Friedman’s test, p<0.05).

## Discussion

### State classification

Although a guideline to standardize sleep staging was published after the discovery of sleep stages (Rechtschaffen & Kales, 1968), staging has remained an inconsistent and often arduous task. Criticisms of and modifications to the guideline (Himanen & Hasan, 2000; Silber et al., 2007) as well as various automated methods (Hamida & Ahmed, 2013; Penzel & Conradt, 2000; Sun et al., 2020) have been developed, but each approach differs in the set of defined states and the type and amount of data required. In addition, studies on sleep typically use electroencephalogram (EEG) LFP recordings, which are distinctly different from intracortical LFPs due to their lower spatial specificity and the complex frequency and phase filtering of bone and tissue, especially at higher frequencies (Buzsáki et al., 2012; Michel et al., 2004; Sejnowski et al., 2014). Beyond these difficulties, manual scoring is also compromised by high inter-scorer variability (Himanen & Hasan, 2000; Younes et al., 2016).

As a result, we developed our own classification methodology to distinguish between different sleep states. The process was relatively simple, utilizing basic dimensionality reduction and clustering on processed LFP PSDs and accelerometer values, but was remarkably consistent. We used several metrics to confirm the method – EOG recordings, concurrent recordings with the Kinect to track movement, k-fold cross validation, and changes in spike rates; they all indicated accurate and robust classification. One explanation may be that sleep states can vary between different regions of the brain (Mascetti, 2016; Siclari & Tononi, 2017). Our focus on a small region (4x4 mm) of the motor cortex and the high spatial specificity of intracortical LFPs may have produced more robust results compared to those based on EEG recordings from across the brain.

Our method could potentially be improved by using a dimensionality reduction method with higher complexity, such as a deep neural network (Malafeev et al., 2018), different clustering methods, such as hierarchical clustering (Gerla et al., 2019), and by including the EOG recordings or other biophysical measurements as additional dimensions of data (Khalighi et al., 2013). However, developing a flawless classification method was not within scope of the study and the method presented was deemed to be sufficient.

### LFP and spike dynamics during different sleep states

Delta band LFP, or slow waves, is typically elevated across the brain during all stages of NREM (Rembado et al., 2017; Silber et al., 2007; Xu et al., 2019). Beta power is high when the animal is awake, particularly in the primary motor cortex as beta oscillations are highly relevant in motor control (Engel & Fries, 2010; Khanna & Carmena, 2017). These two metrics were used as criteria for assigning the classified clusters to the appropriate states. However, the state corresponding to REM sleep, as further confirmed through EOG recordings, presented high theta power, a criterion we did not impose on the classification scheme. The role of theta band in the cortex is still obscure, but theta in the hippocampus is also consistently high during REM (Boyce et al., 2016). We found similar features in the pairwise magnitude-squared coherence; as coherence shows synchrony between two signals, it effectively emphasizes that the increased power is reflective of changes in coordinating activity.

We also observed decreased spiking activity with deeper sleep, similar to previous studies (Steriade et al., 2001; Xu et al., 2019). In addition, the firing patterns of spikes changed due to sleep state – RS neurons had increased high frequency activity during NREM compared to Move, and all neurons had an increase low frequency activity during Rest, REM, and NREM compared to Move. The increase in beta frequency activity of neurons during REM and Rest can be explained by the fact that beta is seen as a “resting” rhythm in the primary motor cortex (Engel & Fries, 2010). RS neurons are likely to be excitatory pyramidal cells and FS neurons are likely to be inhibitory interneurons (McCormick et al., 1985), which may explain the higher “bursty” firing pattern of RS neurons during NREM sleep, potentially due to reactivation of cortical circuitry activated during the day.

### Cross-frequency coupling and spike-LFP synchrony

Cross frequency coupling shows that the higher frequency band modulates with the lower frequency band. Such synchrony reflects coordination of local networks operating on shorter time scales to distributed circuits synchronized at longer time scales; this could potentially play a role in neural computation of attention, learning, and memory (Canolty & Knight, 2010; Jensen & Colgin, 2007). Various measures have been proposed to quantify phase-amplitude coupling; compared to other proposed measures the MVL introduced by Canolty et al. (2006) has been shown to be accurate, the most sensitive to modulations in coupling strengths, and ideal for high quality signals over a long recording epochs (Canolty et al., 2006; Hülsemann et al., 2019; Onslow et al., 2011; Tort et al., 2010).

Spike-LFP synchrony provides another measure of synchrony between the single neurons and the composite synaptic activity of the local population (Buzsáki et al., 2012; Murthy & Fetz, 1996b; Okun et al., 2010). The magnitude indicates the strength of synchrony whereas the phase can provide insights into the timing of the spikes relative to the local population.

Most instances of cross-frequency phase-amplitude coupling and spike-LFP synchrony were stronger for deeper sleep states. This may be due to asynchronous activity during Move being driven by functional local circuitry (i.e. generating movement) whereas activity during resting and sleep states are more attuned to baseline macroscopic rhythmic activity, potentially related to homeostatic plasticity (Tononi & Cirelli, 2014).

#### Delta and high gamma

One noticeable deviation from deeper sleep resulting in stronger synchrony was delta phase to high gamma amplitude (delta-high gamma) coupling, which was significantly higher during Move compared to Rest or REM. Delta oscillations may arise in the thalamus or the cortex and are elevated during NREM. Delta has also been linked to various perceptual, sensorimotor, and cognitive operations, potentially suppressing activity not necessary to the task at hand (Güntekin & Başar, 2016; Harmony, 2013). In contrast, high gamma band LFP is representative of local spiking activity, often strongly correlated with action potentials, and can also reflect neuronal synchrony (Ray et al., 2008; Ray & Maunsell, 2011).

Evidence suggests the strength of delta-high gamma coupling is modulated by dopamine in both the prefrontal cortex and the cortico-basal ganglia network (Andino-Pavlovsky et al., 2017; Güntekin & Başar, 2016). Delta-high gamma coupling has also previously been shown to be present during sleep in both the hippocampus and the neocortex (Clemens et al., 2009; Takeuchi et al., 2015), but the underlying mechanisms are still unclear. Andino-Pavlovsky et al. (2017) showed high delta-high gamma coupling during slow wave sleep in the rodent prefrontal cortex, and Takeuchi et al. (2015) showed strong delta-high gamma coupling during REM and several NREM substages in the primate hippocampus (Andino-Pavlovsky et al., 2017; Takeuchi et al., 2015). However, neither study pursued the state-dependent comparisons, and their broad definition of gamma (>25 Hz) makes independent interpretation difficult. Specific comparisons of delta-high gamma coupling during different sleep states have not been reported.

Spike synchrony to delta band sheds additional light on the delta-high gamma relationship. In our study, only RS neurons showed higher delta-high gamma coupling during Move, and the phase of synchrony was consistently located at the falling edge of the wave (Figure 14). During NREM both RS and FS spikes were synchronized to delta, but the phase of synchrony was immediately after the trough of the wave. These results suggest delta as a coordinating signal that reactivates cortical circuitry during sleep for consolidation (Gulati et al., 2017; Ramanathan et al., 2015; Xu et al., 2019). Spiking activity during the day potentially drives delta activity, as it occurs right before delta is the most depolarized, and delta during NREM may reactivate the neural circuity as it has the highest depolarization right before spiking activity. Similar phase specificity is also reflected in delta-high gamma coupling during NREM.

#### Theta in the motor cortex and REM sleep

Theta oscillations have often been observed during REM sleep, and have also been shown to coordinate hippocampal place cell activity during active exploration (Buzsáki & Moser, 2013; Cantero et al., 2003). In the cortex, modulation of theta has been most prominently seen with theta burst stimulation (TBS) – bursts of gamma frequency stimulation delivered at theta intervals, typically with TMS over the motor region. TBS has been shown to increase the excitability of the motor system, observed through increased motor evoked potentials, potentially by mimicking the coupling of theta and gamma oscillations (Cárdenas-Morales et al., 2010; Huang et al., 2005; Suppa et al., 2016). Additional evidence shows that strengthening theta-high gamma coupling with transcranial alternating current stimulation (tACS) increases motor learning (Akkad et al., 2021). In agreement with these previous findings, our results show much higher theta-high gamma coupling during Move.

Theta has also been shown to be coupled to low gamma. A wealth of recent cross-frequency coupling literature has reported theta-low gamma coupling in both hippocampus and the neocortex, showing modulations dependent on task performance, cognitive engagement, and memory formation (Buzsáki & Wang, 2012; Canolty et al., 2006; Lisman & Jensen, 2013). Although stronger theta-low gamma coupling has also been observed during REM sleep, it was absent in human studies, potentially due to hippocampal theta oscillations in humans being slower than those in rodents (Cantero et al., 2003; Clemens et al., 2009; Jacobs, 2014). Similar to the human studies, we did not find significant theta-low gamma coupling during REM, but we also did not find significant delta-low gamma coupling.

Instead, we found very high theta-beta coupling during REM sleep. Theta-beta coupling is not a common topic of study; there is some evidence that suggests it plays a role in working memory and decision making, but the literature is sparse and focuses on the frontal lobe (Axmacher et al., 2010; Cohen et al., 2009; Liang et al., 2021). In our results, the maximum beta amplitude occurs right before the peak of theta (Figure 9), which means minimum beta occurs right after the trough of theta. If beta is a regulating, inhibitory rhythm in the motor cortex (Engel & Fries, 2010; Khanna & Carmena, 2017; Kilavik et al., 2013), perhaps theta is increasing the excitability of the cortex by minimizing beta, thus inducing disinhibition. This may lead to more effective memory consolidation often seen during REM sleep, relevant to motor skill learning. However, more research is needed to confirm these speculations.

### Concluding Comments

Our study provides the first comprehensive analysis of cross-frequency phase amplitude coupling and spike-field synchrony across all frequency bands within the cortex during different behavioral states. We observed an increase in burst firing and high coordination between spikes during NREM, consistent with previous findings suggesting reactivations of cortical circuity during NREM. High cross-frequency phase-amplitude coupling between delta and high gamma when the animal is awake and moving and during non-REM sleep, as well as spike-field synchrony with delta band LFP, suggest that delta may encode and subsequently drive these reactivations. Modulations in theta to low gamma phase-amplitude coupling, commonly observed in rodents, was not observed, similar to previous human studies. Instead, we observed high theta to beta coupling during REM, potentially driving motor learning by increasing the excitability of the cortex. These results support previous findings and may serve as the basis for future studies into the roles of LFP frequency bands and different sleep states.

## Acknowledgements

We thank Larry Shupe for programming and software support and Rebekah Schaefer and Becky Burch for assistance with animal care, handling, training, and surgery. This work was supported by the National Institutes of Health (NS012542, RR00166, and NS118781), the National Science Foundation (EEC-1028725), and by the National Institutes of Health Office of the Director, Office of Research Infrastructure Programs (ORIP) under award number P51OD010425 and U42OD011123. The authors declare no competing financial interests.

